# Reduced vomeronasal receptor diversity and impaired pheromone detection in Tex15 knockout mice

**DOI:** 10.64898/2025.12.03.690614

**Authors:** Nader Boutros Ghali, Paige Kramer, Joshua Danoff, Ryan Edwards, Hetanshi Patel, Nusrath Yusuf, Zain Zaidi, Kevin Monahan

**Author notes:** Correspondence to be sent to Kevin Monahan. This study conforms to all the relevant regulatory and ethical requirements set forth by Rutgers University IACUC.

## Abstract

The mouse vomeronasal organ (VNO) detects pheromones, which provide vital information needed to find a mate, detect predators, or alert to the presence of an intruder. Within the VNO, pheromones are sensed by two families of G-protein Coupled Receptors (GPCRs) the type 1 and type 2 vomeronasal receptors (V1Rs and V2Rs). However, little is known about the regulatory mechanisms that control the expression of V1R and V2R genes by vomeronasal sensory neurons (VSNs). Here we identify a transcriptional repressor, *Testis-expressed gene 15* (*Tex15*), that regulates vomeronasal receptor gene choice and is required for the proper functioning of the vomeronasal system in regulating male behavior. We find that Tex15 is transiently expressed by VSN precursors prior to strong VR gene expression. The absence of *Tex15* results in a pervasive change in the expression of V1R and V2R genes, which results in a less diverse repertoire of VR expressing cells and includes a dramatic reduction in the expression of specific receptors that have been tied to intermale aggression. Accordingly, *Tex15* knockout mice exhibit lowered activation in the Accessory Olfactory Bulb (AOB) after exposure to male odorants, and a loss of stereotyped aggression between male mice. Taken together, these results show that *Tex15* plays a critical role in pheromone sensing by ensuring that VSNs express a diverse set of receptor proteins.

## Introduction

Olfaction plays a critical role in the survival and reproductive success of many terrestrial animals. The vomeronasal organ (VNO) is the key organ for detecting conspecific and allospecific pheromones from the environment, which can convey vital information such as another animal’s social position in the hierarchy, the receptivity of a female to mating, and the presence of unfamiliar intruder animals (Fu et al., 2015; Ibarra-Soria et al., 2014; Itakura et al., 2022; Leypold et al., 2002; Wysocki, 1979). Within the VNO, vomeronasal sensory neurons (VSNs) are responsible for detecting chemical signals and transmitting this information to the anterior olfactory bulb (AOB), from which it is relayed further on to the relevant neuronal circuits (Itakura et al., 2022). Mice with mutations that affect VSN function exhibit impaired social behaviour such as a lack of aggression in males or abnormal sexual behaviour in females (Ibarra-Soria et al., 2014; Itakura et al., 2022; Kondo et al., 2024; Leypold et al., 2002).

In mice, VSNs recognize chemical signals using G-protein coupled receptor proteins from the type 1 and type 2 families of vomeronasal receptors, known as the V1Rs and the V2Rs. The V1R genes are expressed in a monogenic and monoallelic fashion by VSNs in the apical layer of the epithelium, such that each apical VSN expresses only one type of receptor out of the 230 genes in the V1R family (Dietschi et al., 2022; Hills et al., 2024). In contrast, basal layer VSNs express specific combinations of the 160 V2R genes. The V2R genes are split between four sub-families: A, B, C, and D. Each basal VSN expresses one A, B, or D family gene together with one or more genes from the C subfamily, which includes seven members: *Vmn2r1-7*. The pairing is not random, as individual A, B, D genes tend to be expressed together with either *Vmn2r1* or with the other C family members. Expression of A, B, and D genes precedes expression of C genes, so the choice of an A, B, or D gene may instruct subsequent C family expression (Akiyoshi et al., 2018; Ishii & Mombaerts, 2011). In addition, in rodents a small number of VSNs express a receptor from a family of immune GPCRs, the formyl peptide receptors (Fprs). This family expanded in the rodent lineage after an ancestral Fpr was translocated into a VR gene cluster and gained expression in VSNs (Boillat et al., 2021; Dietschi et al., 2022).

VSN progenitors commit to expressing a type 1 or type 2 VR gene over VSN differentiation. VSNs originate from a population of globose basal cells (GBCs) found in the marginal zones of the VNO (Dietz et al., 2025). These GBCs differentiate into immature VSNs (iVSNs) that are committed to either the apical (V1R-expressing) or basal (V2R-expressing) lineage (Dietz et al., 2025; Hills et al., 2024). The mechanism behind this divergence involves multiple regulatory proteins, with differentiating basal cells expressing combinations of ATF5, Bcl11b, and Notch signaling factors (Katreddi et al., 2022). Perturbing these regulatory proteins results in a disorganised VNO with a different distribution of apical and basal VSNs and a corresponding change in VR expression (Katreddi et al., 2022; Lin et al., 2018; Nakano et al., 2015). However, beyond the specification of neuronal type, the mechanism that controls the choice of VR genes for expression remains unknown.

Here we show that a transcriptional repressor known for its role in repressing transposable elements in the male germ-line, *testis expressed gene 15 (Tex15)*, is expressed by iVSNs and is required for wild-type patterns of VR gene expression, VNO function, and stereotyped intermale aggression. We show that differentiating VSN progenitors transiently express *Tex15* RNA and protein. The peak of *Tex15* transcription coincides with the split between apical and basal VSNs lineages. Mice that are homozygous for a *Tex15* null allele exhibit altered expression of VR genes and altered abundances of neurons expressing an individual VR gene. When exposed to soiled male nesting material, *Tex15* null mice exhibit reduced c-Fos expression, a maker of neuronal activity, in downstream neurons in the accessory olfactory bulb. Finally, male *Tex15* null mice fail to attack intruder males in a resident-intruder paradigm and instead exhibit increased exploratory anogenital sniffing behaviour.

## Results

Single-cell RNA-seq has established *Tex15* as a marker of the immediate neuronal progenitor stage in the olfactory sensory neuron lineage in the main olfactory epithelium (Brann et al., 2025; Pourmorady et al., 2024), but its expression in the VSN lineage has not been characterized. To determine whether *Tex15* is expressed at any point during VSN differentiation, we examined previously published single-cell RNAseq data from mouse VNO (Hills et al., 2024). This dataset contains eight mice, four collected at postnatal day 56 (P56) and four collected at postnatal day 14 (P14). Since we were interested in examining the immature VSNs, we used the P14 samples, which have a higher count of immature VSNs but otherwise resemble the P56 samples. This cohort has two males and two females. Consistent with prior work, UMAP analysis generates distinct clusters of cells expressing markers of mature apical and basal VSNs as well as clusters corresponding to various stages of VSN progenitors (Figure S1A). We excluded a cluster of atypical VSNs, noted by Hills et al, that express markers of both apical and basal VSNs, since these cells do not express *Tex15* and their connection to the canonical set of VSN progenitors is uncertain.

*Tex15* expression is restricted to cells in the neuronal lineage (Figure S1B, C). Thus, to examine *Tex15* expression over VSN differentiation, we used marker genes to identify cell clusters corresponding to the neuronal lineage and then isolated and reclustered these cells. We then assigned each cluster to a stage of differentiation based upon the expression of known markers, namely: *Ascl1* for early progenitors, *Neurod1* for immediate neuronal progenitors, *Gap43* for immature VSNs, *Gnai2* for mature apical VSNs, and *Gnao1* for mature basal VSNs (Figure 1A). Notably, *Tex15* transcript levels peaks in the INP population coinciding with the divergence of INP cells into the apical and basal differentiation lineages (Figure 1B, C).

**Figure 1.**
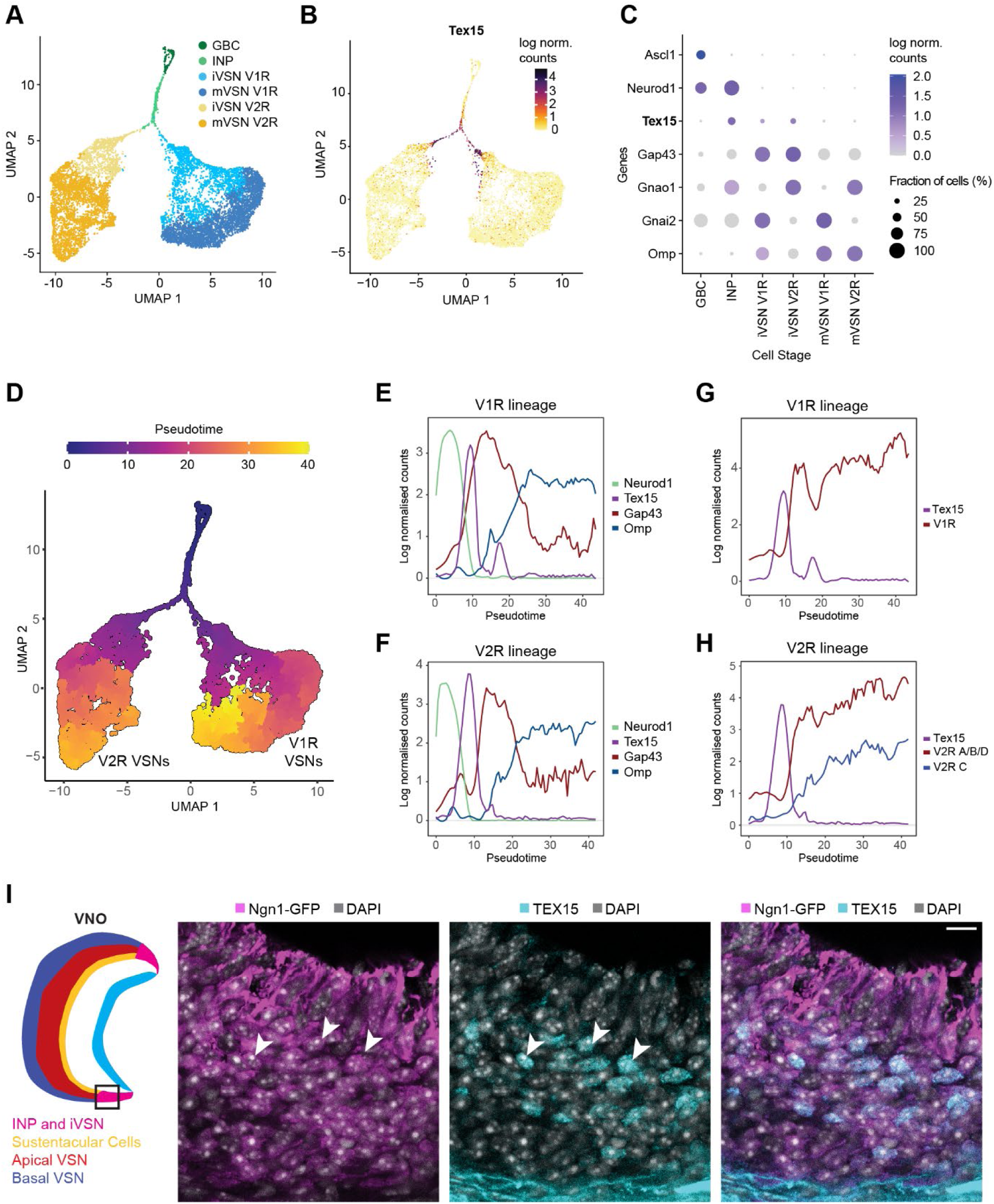
*Tex15* is transiently expressed by VSN progenitors before VR choice. (A) UMAP projection of cells from the neuronal VSN lineage, color coded by stage of differentiation. (B) *Tex15* transcript levels across the neuronal lineage. (C) Transcript levels and frequency of expression for developmental marker genes and *Tex15* in each cell population. (D) Pseudotime values calculated across the neuronal lineage. (E) Transcript levels of developmental markers and Tex15 over pseudotime in the isolated V1R lineage. (F) Transcript levels of developmental markers and Tex15 over pseudotime in the isolated V2R lineage. (G) Transcript levels of Tex15 and the transcript level of the most abundant V1R transcript level in each cell over pseudotime in the V1R lineage (H) Transcript levels of Tex15 and the transcript level of the most abundant V2R A/B/D and V2R C in each cell in the V2R lineage. (I) Immunohistochemistry for TEX15 in VNO tissue sections from postnatal day eight (P8) mice. Images show the marginal zone of the VNO. Scalebar: 10μm. In all plots, transcript levels are expressed as log normalized counts.

To examine the precise timing of *Tex15* expression, we conducted pseudotime analysis of the neuronal lineage (Figure 1D). Consistent with the cell stage analysis, *Tex15* transcripts peak after *Neurod1* and prior to *Gap43* (Figure 1E, F) in both the V1R and V2R lineages. Next, we examined how *Tex15* aligns with the onset VR gene expression. To accomplish this, we determined the highest expressed V1R and V2R genes within each cell in the corresponding lineage. In the basal lineage, we took the additional step of analyzing the C-subfamily V2Rs separately from the A, B, D subfamily V2Rs. We find that in the apical lineage, *Tex15* expression peaks just prior to a dramatic rise in V1R gene transcripts (Figure 1G). Similarly, in the basal lineage, we observed that *Tex15* expression precedes A, B, D subfamily expression, which in turn precedes, C-subfamily expression (Figure 1H). The temporal separation between the rise in A, B, D transcript levels and the rise in C-subfamily transcript levels is consistent with a prior study that observed that expression of A, B and D subfamilies of V2R genes was detectable by RNA in situ hybridisation (ISH) in immature cells prior to the expression of a C subfamily gene (Ishii & Mombaerts, 2011). We conclude that *Tex15* expression is tightly restricted to a time point in the development of VSNs, which precedes VR expression.

We next sought to determine whether TEX15 protein is present in immature VSNs in the VNO. For these experiments, we used VNO tissue sections postnatal day 8 (P8) mice bearing a Neurog1-GFP reporter gene (Gong, et. al., 2003), which labels INPs and iVSNs in the marginal zones. To visualize TEX15 expression, we raised a mouse monoclonal antibody against an epitope in the central region of TEX15 (Figure S2A), and then used it for immunohistochemistry with a GFP antibody to enhance the endogenous Neurog1-GFP signal. We observe *Tex15* immunoreactivity in the nuclei of cells in the marginal zones of the VNO, all of which also exhibit labeling with Neurog1-GFP (Figure 1I). We also observe immunoreactivity along the basal membrane below the basal VSN, but this is retained in Tex15 knockout tissue and thus is likely non-specific (see below). Thus, we conclude that TEX15 protein is abundant in the nuclei of VSN progenitors found in the marginal zone of the VNO.

To examine the effects of *Tex15* we used a *Tex15* null allele (Tex15KO) in which the majority of the *Tex15* coding sequence has been replaced by beta galactosidase (Yang et al., 2008). Homozygous Tex15KO mice and littermate controls were obtained by crossing Tex15 +/-heterozygous mice. To confirm that the knockout lacks *Tex15* expression in the VNO, we performed a *Tex15* immunofluorescence staining in postnatal day 2 KO mice and littermate controls (Figure 2A). As above, nuclear *Tex15* immunoreactivity is present in the marginal regions of the VNO, although we also observe occasional labeled nuclei throughout the rest of the VNO in heterozygous and wildtype animals (Figure S2B). The nuclear signal is completely absent from both regions in Tex15KO tissue. Otherwise, the overall structure and organization of the VNO appear similar in Tex15KO mice.

**Figure 2.**
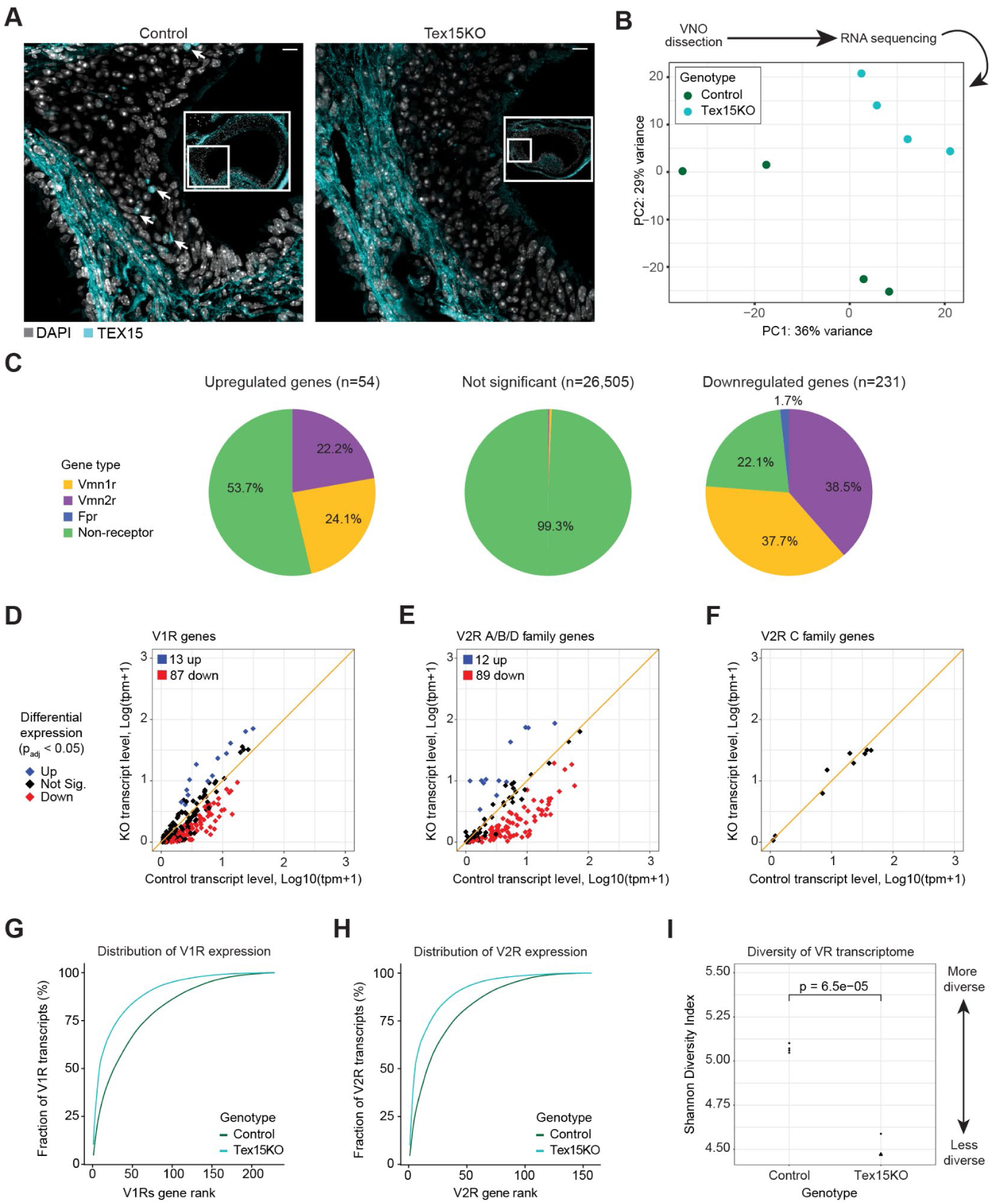
*Tex15* knockout mice have altered VR expression and reduced diversity. (A) Coronal VNO section from a P2 Tex15KO mouse and littermate control labeled with TEX15 antibodies (cyan) and DAPI, which labels DNA (gray). Tex15 immunoreactivity is detected in nuclei of the control mice (arrows). Inset: Stitched tile scan of the full VNO. Scalebar: 10μm. (B) PCA analysis of VNO bulk-RNAseq libraries. (C) Abundance of *Vmn1r*, *Vmn2r*, and *Fpr* genes within the sets of significantly upregulated and significantly downregulated genes in the Tex15KO. (D) Scatter plot of V1R transcript levels in Tex15KO versus control VNOs. (E) Scatter plot of V2R A/B/D family transcript levels in Tex15KO versus control VNOs. (F) Scatter plot of V2R C-family transcript levels in Tex15KO versus control VNOs. (G) The cumulative expression of V1Rs ranked by expression within each genotype. (H) The cumulative expression of V2R A, B, and D family genes ranked by expression within each genotype. (I) Shannon’s Diversity Index of all VR transcripts in control and Tex15KO libraries df = 3.9187, p-value = 6.491e-05 (t-test). For bulk RNA-seq, differentially expressed genes were determined based upon padj < 0.05 for a >50% change in expression (Wald test).

Based upon its expression in INP cells, we hypothesized that Tex15 might regulate the expression of VR genes. To test this, we performed RNAseq on VNO tissue obtained from adult Tex15KO mice and heterozygous controls. We sequenced poly-A selected RNAseq libraries from four animals per genotype, and then performed differential gene expression analysis using DESeq2 (Love et al., 2014). As expected, the samples cluster by genotype when subjected to principal component analysis (Figure 2B). Differential expression analysis identifies 283 genes that exhibit a greater than 50% change in expression between KO and Het animals (p_adj_ < 0.05 for >50% change in expression) Most differentially expressed genes are V1Rs and V2Rs: 151 out of a total of 283. However, the total expression of V1R and V2R genes is not affected; the sum of transcript counts from each lineage are similar between Het and KO animals (Figure S3A). Rather, the expression of individual receptors is significantly different (Figure 2D). Notably, the changes to V2R expression are restricted to the A, B, and D subfamilies, 102 out of 160 genes of which are differentially expressed in the KO (Figure 2E). None of the seven C-family V2Rs, which are co-expressed depending on prior A, B, or D choice, are differentially expressed in the Tex15KO (Figure 2F). Interestingly, we also see reduced expression of more than half of the FPR genes, with 4 significantly decreased in Tex15KO mice (Figure S3B).

Taken together, these results suggest that Tex15 regulates the choice of a receptor protein by VSNs, but suggests that it may not affect the downstream processes that influence the subsequent selection of a C-subfamily V2R. Consistent with this idea, among the non-VR genes that are differentially expressed in the Tex15KO VNOs are four H2-MV group genes: *H2-M1*, *H2-M9*, *H2-M11*, and *H2-M10.4* (Figure S3C). The H2-MV genes are expressed in specific combinations with VR genes, and the first three of these are specifically co-expressed with *Vmn2r82* and *Vmn2r81 (Devakinandan et al., 2024)*, which are downregulated in the knockout (Figure S3D).

Overall, the Tex15KO results in higher expression of a few VR genes, whereas many more exhibit decreased transcript levels. This results in a reduction in the diversity of the VR gene repertoire (Figure 2G, H). We quantified VR gene diversity by calculating the Shannon diversity index value for all VR gene expression levels in Tex15KO and control samples. We observe a significant reduction in these scores, indicating reduced VR diversity in KO mice (t(df) = 3.9187, p-value = 6.491e-05) (Figure 2I).

VR and FPR genes are arranged in clusters across chromosomes, and this clustered organization contributes to their regulation (Dietschi et al., 2022). We asked whether the altered expression of VR genes in Tex15KO animals reflected the chromosome cluster location of each VR gene. Grouping VR genes by cluster did not reveal any consistent patterns as to which VRs were upregulated and which were downregulated. Many VR clusters contain both upregulated and downregulated VR genes, nor are there any individual VR clusters that exhibit uniform increase or decrease in expression (Figure S3E, F).

Since V1R and V2R A, B, and D genes are expressed monogenically within the population of VSN, the changes we observed by bulk RNAseq could arise from either a change in the number of cells expressing each VR genes, reflecting a change in the frequency of VR choice, or a change in the levels of VR expression within the same number of cells. To test whether differentially expressed VR genes are expressed by a different number of cells in Tex15KO animals, we performed RNA in situ hybridization (ISH) to label individual cells expressing specific VR genes in tissue sections from the VNO. Due to the very high sequence similarity among VR genes, we relied on a chromogenic ISH strategy (BaseScope) that can target genes that vary over only short regions, as few as 50 base pairs. For this analysis, we first sought to identify candidate VR genes that had large differences in raw expression values (transcripts per million) as well as a large fold change between conditions. Even with a 50 base sequence length there was only one VR which met our criteria and was not predicted to cross react with other VR genes, *Vmn1r202* (log2fold change = −3.45, p_adj_= 4.89e-35). As a positive control for the BaseScope assay, we also examined the expression of a C family VR, *Vmn2r5*, that is not differentially expressed (log2fold change = 0.09, p_adj_= 0.20). We dissected the VNOs of Tex15KO and littermate controls and counted the nuclei present in the epithelial layer to see what percentage of nuclei were labeled by each probe. As expected, the *Vmn2r5* probe is widely expressed in the basal layer of the VNO (Figure S4A) and is similar in Tex15KO and control tissue sections. In contrast, we observe significantly fewer cells expressing *Vmn1r202* in Tex15KO mice (F(1,3.63)=11.68, p=0.031, using a mixed linear model with a fixed effect of genotype and a random effect of subject), in accordance with the transcriptional changes observed by bulk RNA-seq of Tex15KO and control VNO (Figure 3A). Importantly, we still observe strongly positive cells in the KO animals (Figure S4B), but at a reduced rate compared to control tissue, which is consistent with an alteration in the frequency of receptor choice rather than expression per cell.

**Figure 3.**
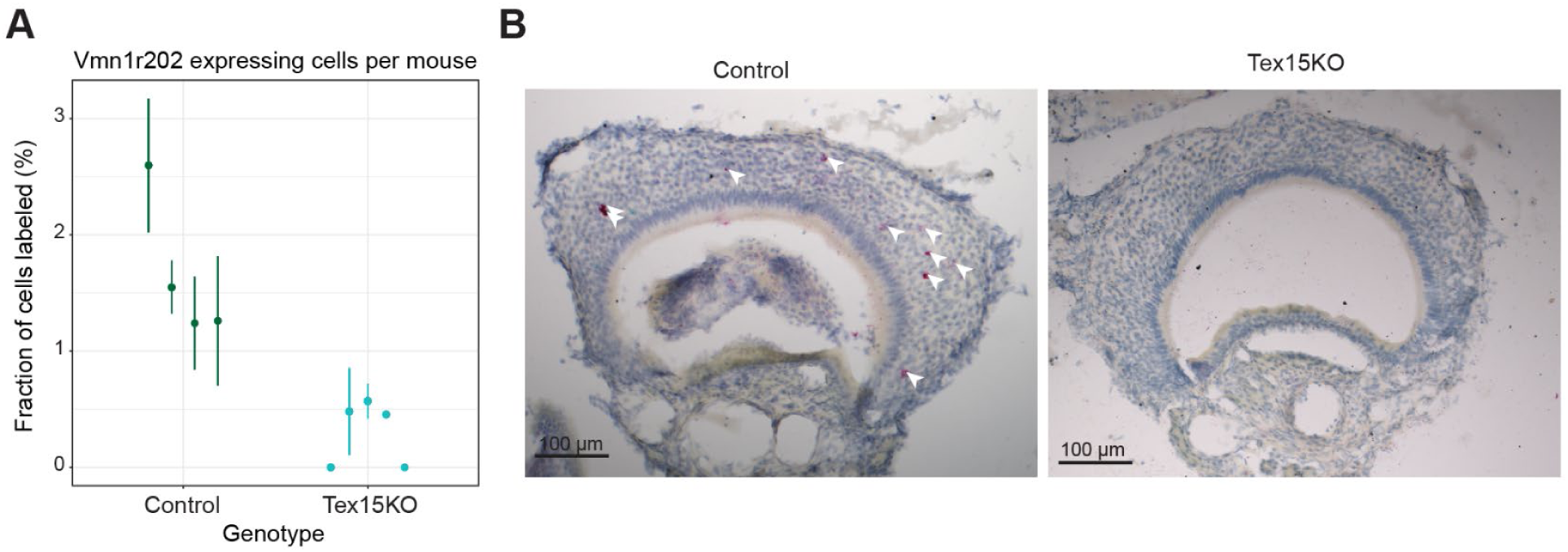
Tex15KO results in fewer cells expressing *Vmn1r202*. (A) The percentage of cells within the VSN layer expressing *Vmn1r202* in Tex15 knockout vs control (p=0.006). Each point-line represents the mean and standard error determined by counting multiple sections per mouse. (B) Representative images of a knockout and control VNO. *Vmn1r202* BaseScope chromogenic signal is red. Scalebar: 100μm.

Our prior results indicate that Tex15KO VNOs contain a reduced diversity of VSNs. Since the expression of a diverse repertoire of VRs is thought to facilitate the detection of signals from conspecifics, we asked whether the reduced VSN diversity in Tex15KO mice might impair the response to mouse derived odorants. We tested this hypothesis by quantifying the expression of c-Fos, an immediate early gene that is expressed in response to neuronal activity in the mitral cells of the accessory olfactory bulb (AOB). These are the first layer of cells downstream of VSN and serve to process the information encoded in pheromones and relay it to the downstream targets in the brain (Imamura et al., 2020). To test this, *Tex15* knockout mice and wildtype controls were placed in a new cage that contained either soiled male nesting material or clean nesting material for 2 hours (Figure 4A), and then we assayed for c-Fos expression in the AOB. To facilitate identification of the AOB, these experiments were performed with mice bearing an OMP-ires-GFP reporter gene, which labels VSN axons located within the glomeruli of the Accessory Olfactory Bulb (AOB) (Figure 4B). As expected, there is a significant genotype by condition interaction (F_(1,8)_=6.64, p=0.033) (Figure 4C). Wild type mice exposed to dirty bedding had significantly higher rates (p = 0.0121) of c-Fos positive nuclei (32.3%± 2.63%) than wildtype mice exposed to clean bedding (20.3% ± 3.79%). In contrast, *Tex15* knockout mice exposed to clean bedding and dirty bedding have similar rates of c-Fos expression (17% ± 2.95% vs 18.7% ± 2.36%, p = 0.5697). Moreover, c-Fos expression in the AOBs of Tex15KO mice exposed to dirty bedding was similar to c-Fos express in the AOBs of wildtype mice exposed to clean bedding (p= .9357), supporting a failure to detect male pheromones in the KO. Taken together, these results suggest that the alterations in VR expression rates observed in Tex15KO mice impair the response of the vomeronasal system to male derived odorants.

**Figure 4.**
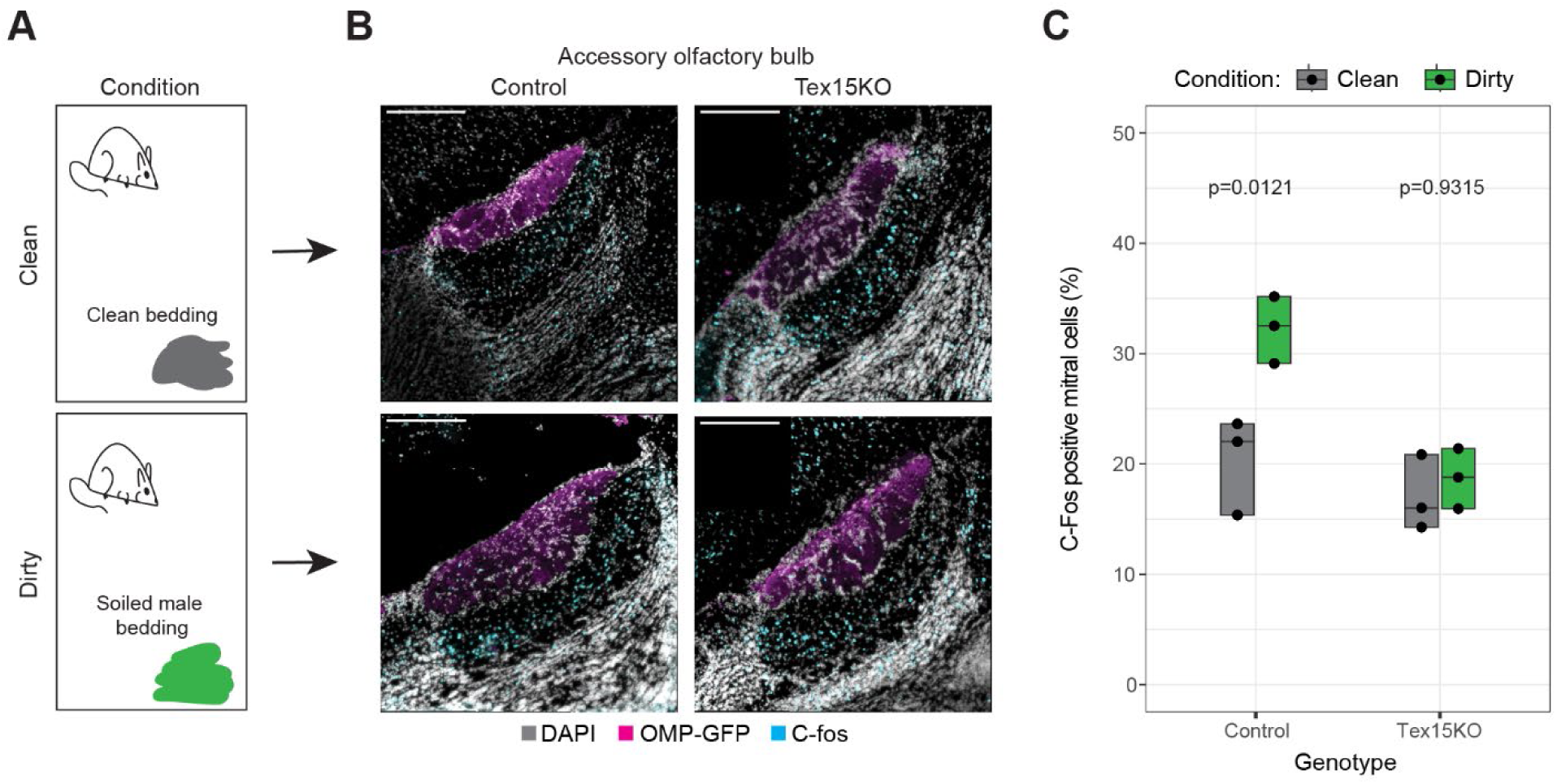
Loss of *Tex15* and VR diversity results in lower activation in the AOB. (A) Schematic showing the soiled bedding exposure assay. (B) Representative immunohistochemistry images showing c-Fos and GFP signal in sagittal sections of the accessory olfactory bulb. The images shown are tile scans that were stitched together. Scalebar: 250μm (C) The average percentage of activated neurons per mouse, grouped by genotype and condition. Mean values for each mouse were calculated by averaging across 3 sections. Statistical significance was determined using a mixed linear model, which included a fixed effect of genotype, condition, and the interaction of genotype and condition, and a random effect of subject.

The impaired activation of the AOB in response to male odorants suggests that Tex15KO mice may also exhibit altered behavior in response to male cues. The VNO plays a critical role in many innate behaviours, such as intermale aggression or identifying the sex of the other mouse (Leypold et al., 2002). To test VNO function in Tex15KO mice we chose the resident intruder experiment, as a functioning VNO is key to eliciting the stereotyped aggression from the male resident (Itakura et al., 2022; Leypold et al., 2002). A male mouse in an established territory will react aggressively toward an unknown male intruder (Leypold et al., 2002). This assay typically uses sexually experienced males as the resident, as they exhibit greater responses than sexually naive males (Koolhaas et al., 2013). However, due to widespread changes in VR receptor diversity, we chose to use sexually naive males for these experiments, to avoid potentially confounding effects of *Tex15* on mating behavior.

We ran the resident intruder with three 10-minute trials over the course of three days while comparing control and *Tex15* knockout naive males interacting with intact intruders that are age-matched (Figure 5A). An attack latency of 600 seconds indicates that the resident did not initiate an attack during the trial. However, Tex15KO males exhibited significantly different behaviour as calculated by Fisher test (p= 0.002997); Tex15KO male residents never initiated an attack against an intruder at any point during any of the trials (Figure 5B). We noted that the knockout residents spent more time anogenitally sniffing their intruders when compared to the control (p=0.048) (Figure 5C).

**Figure 5.**
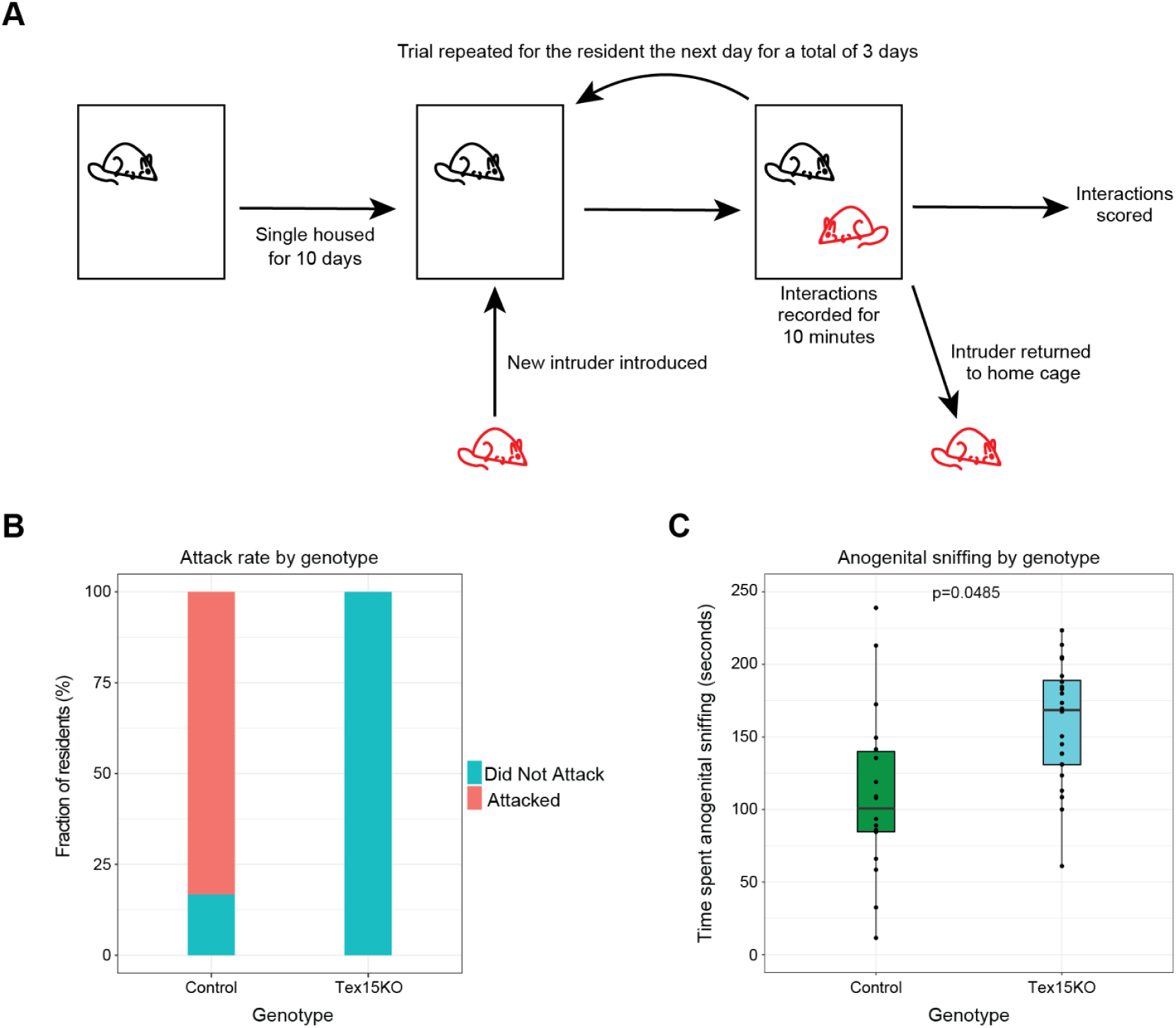
*Tex15* knockout males do not show aggressive behaviour towards intruders. (A) Diagram of the resident intruder assay conducted on the mice. (B) Percent of trials in which the resident attacked the intruder. (C) Plot of the amount of time control and Tex15KO mice spent in each trial performing anogenital sniffing behavior.

## Discussion

We find that *Tex15* is expressed at a critical time point in VSN development, coinciding with the divergence between the basal and apical populations and shortly preceding the onset of VR gene transcription. In mice lacking *Tex15*, the overall expression of VR genes is unchanged, but the relative abundance of individual VR genes is altered. Most VR genes exhibit reduced expression, whereas a few are expressed at higher levels, resulting in a reduction in the overall diversity of VR gene expression. We show that the downregulated genes are a result of lowered frequency of being chosen. These transcriptional changes are accompanied by reduced AOB activation in response to soiled male nesting material. We also observe that Tex15KO mice exhibit a lack of stereotyped intermale aggression and increased ano-genital sniffing behaviour. Taken together, these findings indicate that *Tex15* has a crucial role in ensuring that VR receptor choice results in diverse outcomes, generating a VNO in which many subtypes of VSN are represented.

The transient expression of *Tex15* during a precise developmental window, coupled with the striking alterations in VR gene expression observed in *Tex15* knockout mice, implicates Tex15 as an important regulator of VR gene choice. The loss of *Tex15* affects all of the main chemoreceptor receptor families expressed by VSNs, the V1Rs, the subfamily A, B, and D V2Rs, and the FPRs, implying these receptors share a *Tex15*-dependent choice mechanism. Conversely, gene regulatory decisions that are downstream of VR choice, such as V2R C family expression and H2-MV expression, seem to respond to the overall changes in the VR repertoire.

What then is the role of *Tex15* in regulating VR gene choice? In testes, *Tex15* associates with PIWI family proteins MILI and MIWI2, which mediate piRNA-directed methylation and silencing of transposable elements. Tex15KO mice exhibit reduced DNA methylation of LINE1 and LTR family transposable elements in testes, which is accompanied by increased transposon expression (Schöpp et al., 2020; Wang et al., 2005; Yang et al., 2020). Expression of PIWI family members is very low or absent in the VNO, making it unlikely that the altered VR expression results from altered piRNA directed gene regulation. However, it is possible that *Tex15* associates with the same gene silencing machinery that carries out the piRNA directed DNA methylation and heterochromatin deposition on transposable elements. This would implicate a role for heterochromatin deposition in regulating VR gene choice, similar to what has been proposed for the olfactory receptor genes (Bashkirova et al., 2023; Magklara et al., 2011). These results also suggest a possible role for small RNA directed silencing in VR gene regulation, which has an incompletely understood role in the olfactory system (D. Yang et al., 2020). Identifying the protein partners of Tex15 in the VNO will be critical for characterizing its tissue specific mechanism of action.

Alternatively, it is possible that the observed effects on VR gene expression are a direct result of dysregulation of transposable elements in Tex15KO mice. The transcription start sites (TSSs) of over half of VR genes were found to be upstream of LINE1 elements and, moreover, these LINE1 elements are transcribed (Pascarella et al., 2014). It may be that this LINE1 transcription influences subsequent VR gene choice, and that alterations in this transcription in Tex15KO mice results in the observed changes in VR gene expression. In this scenario, variation in the proximity of transposable elements to the transcriptional start site of VR genes may account for the variable effect of the Tex15KO on VR gene expression. Future work should examine the Tex15KO mice for potential changes in transposable element regulation that might be upstream of altered VR expression.

We observe striking defects in the function of the vomeronasal system in Tex15KO mice. Indeed, the responses to male bedding observed in the accessory olfactory bulb of Tex15KO mice are comparable to those from wildtype mice housed in a clean environment. It is unclear whether these differences arise directly from the altered repertoire of expressed VR genes, or some other defect in VSN signaling in Tex15KO mice. However, this loss of sensitivity to male odorants aligns with a dramatic reduction in aggressive behavior in Tex15KO male mice in a resident-intruder behavioural assay. Tex15KO male mice never attacked a male intruder across any of our trials. This loss of aggressive behaviour was accompanied by a significant increase in anno-genital sniffing behaviour. We speculate that this increased exploratory behaviour may reflect a response to deficient signaling through the VNO pathway.

We show that *Tex15* expression is restricted to a key stage of VSN differentiation, corresponding to the split between the V1R and V2R lineages, and directly preceding the onset - of VR gene expression. As such, *Tex15* could serve as an excellent marker for this crucial stage of differentiation. A large portion of the literature has focused on this stage of VSN differentiation (Devakinandan et al., 2024; Enomoto et al., 2011; Katreddi et al., 2022; Lin et al., 2018). As such, Tex15 may serve as a useful driver for genetic tools designed to target this population. Thus, our work identifies *Tex15* as both a key regulator of VR choice and VNO function as well as a useful marker for a key stage in VSN differentiation.

## Methods

| REAGENT or RESOURCE | SOURCE | IDENTIFIER |
| --- | --- | --- |
| **Antibodies** |  |  |
| C-Fos Antibody | Abcam | Cat#ab190289 |
| GFP Antibody | Antibodies Inc | Cat#gfp-1020 |
| 1H11-A3-5 Tex15 Antibody | Homemade |  |
| IgY (H+L) Goat anti-Chicken, Alexa Fluor® 488, Invitrogen | Fisher | Cat#A11039 |
| Cy™3 AffiniPure Goat Anti-Mouse IgG, Fcγ subclass 1 specific | Jackson ImmunoResearch | Cat#115-165-205 |
| IgG (H+L) Highly Cross-Adsorbed Donkey anti-Rabbit, Alexa Fluor® 555, Invitrogen | Fisher | Cat#A31572 |
| **Critical commercial assays** |  |  |
| Basescope Duplex Assay | ACD |  |
| Direct-Zol RNA Miniprep | Zymo Research | Cat#R2050 |
| Bulk RNA-seq | Admera Health |  |
| **Deposited data** |  |  |
| P14 VNO Single Cell Dataset | Hills Jr. et. al. | https://doi.org/10.7554/eLife.97356.3 |
| **Experimental models: Organisms/strains** |  |  |
| Tex15 KO mouse | Yang, et. al., 2008 | https://doi.org/10.1083/jcb.200709057 |
| OMP-IRES-GFP | Mombaerts et al., 1996 | https://www.jax.org/strain/006667 |
| Neurog1-EGFP | Gong, et. al., 2003 | #010618-UCD |
| **Software and algorithms** |  |  |
| R software(4.5.2) | R Core Team 2016 | https://www.r-project.org/ |
| RStudio(2024.12.1.563) | Rstudio Team 2020 | https://rstudio.com/ |
| Tidyverse(2.0.0) | Wickham H, et. al. (2019) | [doi:10.21105/joss.01686](https://doi.org/10.21105/joss.01686) |
| Dplyr(1.2.0) | Wickham H, François R, Henry L, Müller K, Vaughan D (2026) | https://dplyr.tidyverse.org |
| ggplot2(4.0.2) | Wickham H (2016) | https://ggplot2.tidyverse.org |
| rstatix(0.7.3) | Kassambara A (2023) | https://rpkgs.datanovia.com/rstatix/ |
| ggpubr(0.6.3) | Kassambara A (2025) | https://rpkgs.datanovia.com/ggpubr/ |
| DEseq2 (3.22) | Love, M.I., Huber, W. and Anders, S., 2014 | [10.1186/s13059-014-0550-8](https://doi.org/10.1186/s13059-014-0550-8) |
| Monocle3(3) | Cao et al, Nature 2019 | https://cole-trapnell-lab.github.io/monocle3 |
| Seurat(5.4.0) | Hao. Y. et al 2023 | https://cran.r-project.org/web/packages/Seurat/citation.html |
| SeuratObject(5.3.0) | Satija R, et. al. 2023 | https://satijalab.r-universe.dev/SeuratObject/citation |
| Vegan(2.7.3) | Oksanen J et. al. 2025 | https://vegandevs.github.io/vegan/authors.html |
| naniar(1.1.0) | Tierney N, Cook D (2023) | https://www.jstatsoft.org/article/view/v105i07 |
| forcats(1.0.1) | Wickham. H. 2025 | [10.32614/CRAN.package.forcats](https://CRAN.r-project.org/package=forcats) |
| Fiji(2.17.0) | Schindelin et al. 2012<br>DOI: [doi:10.1038/nmeth.2019](https://doi.org/10.1038/nmeth.2019) | https://imagej.net/software/fiji/ |
| afex(1.5.1) | Singmann H. et. al. 2016<br>DOI: [10.32614/CRAN.package.afex](https://CRAN.r-project.org/package=afex) | https://CRAN.R-project.org/package=afex |
| emmeans(2.0.2) | Lenth & Piaskowski 2026 | https://rvlenth.github.io/emmeans/authors.html |
| Trim Galore (0.6.5) | https://doi.org/10.5281/zenodo.5127898 | https://github.com/felixkrueger/trimgalore |
| Cutadapt (5.1) | Martin M. 2011 DOI: 10.14806/ej.17.1.200 | https://cutadapt.readthedocs.io/en/stable/ |
| STAR (2.7.11b) | Dobin et al 2013 | https://github.com/alexdobin/STAR |
|  | DOI:<br>10.1093/bioinformatics/bts635 |  |
| samtools (1.19.2) | Danecek et al 2021 DOI:<br>10.1093/gigascience/giab008 | https://www.htslib.org/ |
| htslib (1.21) | Bonfield et al 2021 DOI:<br>10.1093/gigascience/giab007 | https://www.htslib.org/ |

### Tex15 Antibody Generation

DNA encoding the Tex15 immunogen was cloned from mouse genomic DNA then inserted into pGex4T-2. The protein was expressed in *E. coli* and purified from the insoluble pellet by solubilizing with Sarkosyl, TX100 and Chaps at a ratio of 2% to 4% to 2.5%. Mice were immunized with 100 ug of protein emulsified in Freund’s Adjuvant and were subsequently bled. A mouse was boosted with Tex15 protein in PBS and spleen cells were fused with an NS-1 parental cell line using Roche PEG1500 (Sigma 10783641001). The fused cells were plated in IMDM 20% FCS with HAT selection. Cells were grown for 10 days and then the supernatants from the growing cells were screened for the presence of anti-Tex15 antibody by screening against Tex15 fusion protein in an ELISA. ELISA positives were screened on P5 olfactory epithelia from Tex15KO and heterozygous mice. Clone 1H1 was determined to be positive after screening on tissue blocked with Donkey Fab anti Mouse IgG, 30ug/ml (Jackson ImmunoResearch 715-007-003) for 1 hour at RT or overnight at 4°C prior to staining.

### Tex15 immunogen, aa 597-1070

RDDKNPNEAKEHNTDNINGSEKQDCLANDHFTNIVEMREIKSNTEVEILNSEECFT FNSFRGKNGKPAETASSESEAVEQRHAPNDQRGLEHLVSSFPEIEGSSVCVASNATKQI VGTTVLTVSTSLGDHQKDELKEICSSESSDLGLVKHSISECEIDTDKDKLQDFHQLVN ENSALKTGLGSEIEVDLEHDNASVFQQNMHSQGNDLCEEFELYESLKSRIDWEGLFGS SYEEIESSSFARREGTDQHSSTECNCVSFCSQDKRELHNPIFLPDLQVTITNLLSLRI SPTDESLELKDNFYKQVTESTEPETNKEGNASGFGMCSQPSGENSSFSCANKFGNSVQ ESGDVSKSESSHSSNSSHNTHVDQGSGKPNNDSLSTEPSNVTVMNDKSKCPTKSKPVF NDTRNKKDMQSRSSKRTLHASSSRGQNIANKDLREHETHEKKRRPTSHGSSDRFSSLS QGRIKTFSQS

### Tex15 Immunofluorescence

Mice under P8 were decapitated and their heads were fixed in 4% PFA in PBS for 30 minutes at room temperature. The heads were then washed three times for 5 minutes in PBS before being submerged in a 30% sucrose and PBS mixture at 4°C for 3 days. Afterwards, the heads were embedded in Optimal Cutting Temperature compound (Cat#:25608-930) and then sectioned with a cryostat to a thickness of 14 microns. Sections were transferred onto superfrost plus slides (Cat# 48311-703) and stored at −20C. The slides were washed in a solution of PBS + 0.1% Triton (0.1% PBST) before adding a primary antibody solution of anti-Tex15 mouse monoclonal antibody (1H1A324-3) diluted at 1:10 in 4% goat serum and 0.1% PBST. The slides were incubated with the primary antibody in a humidified chamber at 4°C overnight. The next day, they were washed three times for 5 minutes with 0.1% PBST. The secondary antibody goat anti mouse IgG1 Cy3 (Cat#: 115-165-205) and DAPI (Sigma-Aldrich: MBD0015-1ML) were diluted at 1:1000 in 4% goat serum in 0.1% PBST and applied on the slides. They were incubated in a humidified chamber at room temperature for 1 hour. The slides underwent three more 5-minute washes in 0.1% PBST. They were then mounted with VectaShield Hardset Mounting Medium and left to cure for 15 minutes in the dark. The images were taken with a Leica TCS SP8 tauSted 3X confocal and imaged at 40x. Tilescan images were then stitched together in FIJI with the Grid/Collection Stitch plugin (, et al, 2009).

### Tex15 and Ngn-GFP Immunofluorescence

Tissue sections were prepared as described above, and placed into a blocking solution of 4% goat serum in 0.1% PBST for 1 hour at RT. The slides were then incubated with primary antibodies for Tex15 (1H1A324-3) diluted at 1:10 and anti-GFP (Cat#: gfp-1020) at 1:1000 in 4% goat serum and 0.1% PBST. The slides were incubated in a humidified chamber at 4C overnight. The next day, they were washed three times for 5 minutes with 0.1% PBST. The slides were then incubated with secondary antibodies, goat anti-mouse IgG1 Cy3 (Cat#: 115-165-205), IgY goat anti-chicken Alexa Fluor 488 (Cat#: A11039), and DAPI (Sigma-Aldrich: MBD0015-1ML), diluted at 1:1000 in 4% goat serum 0.1% PBST in a humidified chamber at room temperature for 1 hour. The slides were then washed three more times, each 5-minute with 0.1% PBST, and then were mounted with VectaShield Hardset Mounting Medium and left to cure for 15 minutes in the dark. The images were taken with a Leica TCS SP8 tauSted 3X confocal at 63x. The brightness and contrast of the Tex15 channel was adjusted to reduce non-specific background reactivity.

### RNA extraction and RNA-seq

The VNO was either dissected and used immediately or frozen at −80°C. RNA was extracted by solubilizing the tissue in Trizol (Cat#: 15596026). The extracted RNA was isolated from the Trizol solution using Zymo Research Direct-zol RNA Miniprep Kits (Cat# R2050). Purified RNA was assessed on a Thermo Fisher Scientific Nanodrop One for concentration. It was then quality controlled for RIN in an Agilent Tapestation 4150. Samples which had a RIN greater than 7 were sent to Admera (South Plainfield, NJ, USA) for DNAse treatment, library preparation based upon poly-A tail selection, and paired-end sequencing

### BaseScope Imaging and Counting

BaseScope probes were generated by ACD. Hybridization and chromogenic staining were performed using the ACD fixed/frozen protocol for the BaseScope duplex and BaseScope single channel assay. Images were collected on a Zeiss Axio Imager.A1 with a lumineer camera. Red, green, and blue images were taken and overlayed to create a full colour image.

### C-Fos Imaging and Counting

Mice were placed in clean cages for 10 minutes to habituate. After that time, either clean nesting material or soiled nesting material from male cages was added to the cage, and the mice were incubated for 2 hours with the material. The mice were euthanised with CO2 and their olfactory bulbs were dissected out and fixed on ice for 20 minutes with 4% PFA in PBS. The bulbs underwent two PBS washes on ice before being placed in 50% 0.5M EDTA in PBS at 4°C overnight, followed by a solution of 30% sucrose in PBS at 4°C for 3 days or until the tissue sank to the bottom. The bulbs were embedded sagittally in OCT and sectioned on a cryostat at 14 microns. For each animal, we collected three evenly-spaced 14-micron-thick sections of AOB tissue.

The slides were blocked in a solution of 4% donkey serum and 4% goat serum in 0.1% PBST for 1 hour. They were incubated in the primary antibody solution of c-Fos (Cat#: ab190289) at 1:500 and anti GFP (Cat#: gfp-1020) at 1:1000 to enhance the fluorescence overnight in a humidified chamber at 4°C. The next day, the slides were washed three times for 5 minutes with 0.1% PBST. The secondary antibody donkey anti rabbit Alexa 555, Alexa Fluor 488 (Cat#:A21206), and DAPI (Sigma-Aldrich: MBD0015-1ML) were diluted at 1:1000 in 4% goat and donkey serum in 0.1% PBST and applied onto the slides. They were then incubated in a humidified chamber at room temperature for 1 hour. The slides underwent three more 5-minute washes in 0.1% PBST. The slides were mounted with VectaShield Hardset Mounting Medium and left to cure for 15 minutes in the dark. The brightness and contrast of the c-Fos channel was adjusted to reduce non-specific background reactivity. Images were collected on a Widefield Leica DMI8 Thunder System at 40X. Each image included the glomerular and mitral layer of the AOB.

### Resident Intruder

Resident males were isolated for a period of 10 days prior to the trials. The resident males were naive and had never been housed with a female. A wildtype, naive intruder of similar age and size to the resident would be placed into the resident’s cage and their interactions recorded for 10 minutes and scored for social interactions, anogenital sniffing, and aggression based on how many seconds they spent engaging in each behaviour. The trial was repeated 3 times over the course of 3 days, each time with a new intruder who had not been previously used in a trial.

The recordings were scored by two independent and blinded scorers and an average of the values and scores was generated by determining how many seconds the residents spent engaging in certain behaviours. The scores of knockout residents and wildtype residents were then compared.

### Quantification and Statistical Analysis

Statistical analysis of histology and behavioral data was completed using mixed models to account for repeated measurements from the same subjects. Mixed models were generated, as described below, using the *afex* package in R. Post-hoc analyses were completed using the *emmeans* package in R.

### Single Cell Analysis

Single cell RNA-seq analysis was performed in R using Seurat. Count matrices from GSE252365 were annotated with relevant metadata including age, sex, and replicate number. The replicates were combined into one object and then low quality cells and possible doublets were discarded by filtering on the fraction of mitochondrial reads, total counts, and total RNAs detected. The data was then log normalized, highly variable genes were identified, and these genes were used for principal components analysis. The different samples were then integrated using Seurat’s “integrateLayers” function and Anchor-based Canonical Correlation Analysis. The integrated data was then clustered using 30 PCA dimensions and a clustering resolution of 4. Clusters from the neuronal lineage were identified by examining the expression of established marker genes (Hills et al., 2024), and these cells were selected for further analysis. As noted in Hills et al., 2024, there is a population of VSNs which do not follow the canonically established development trajectory and thus were excluded from the subset.

After subsetting, the cells from the neuronal lineage were reclustered, as described above, and then clusters were identified based on the expression of marker genes. Then a pseudotime analysis was carried out with Monocle3. For subsequent analyses of the V1R VSN and V2R VSN lineages, all cells prior to the lineage split (2 clusters) were retained and then we selectively retained either the V1R VSN or V2R VSN cells, as appropriate. These cells were once again reclustered and used for a lineage specific pseudotime analysis with Monocle3. We then plotted expression over pseudotime by combining pseudotime values for each cell with log normalised gene counts.

### RNA-seq analysis

Adapters were removed from sequencing data using Trim Galore and cutadapt. STAR was used to align trimmed reads to the mouse genome (mm10). For alignment, the lab first used a custom transcriptome annotation file based upon ENSEMBL release GRCm38.102, but updated to include expert curated olfactory receptor and vomeronasal receptor gene annotation from Barnes et al 2020 and Dietschi et al 2022, respectively. Aligned reads were read into R using Rsubread, and then differential expression analysis was performed using DEseq2.

### BaseScope Analysis

The Basescope images were analyzed to determine the number of nuclei that were red positive relative to negative nuclei. The specifications for a positive versus a negative cell were determined by comparing experimental BaseScope images to images that used a negative and control bacterial probe provided by ACD. The BaseScope images were unaltered after acquisition. The signal in the negative images served as a standard for background signal. A positive cell required signal strength higher than the negative control in a concentrated area surrounding or adjacent to a nucleus. FIJI counting plugin was used for counting expressing cells in the epithelial layer of the VNO. Positive cells were divided by the total number of cells to generate a percent positive in each image. Mixed models were used to analyze the number of cells with positive BaseScope signal, with a fixed effect of genotype and a random effect of subject.

### C-Fos Analysis

Each condition included three Tex15KO and three wildtype controls, a total of n=12. The brightness and contrast of the c-Fos and DAPI channels were adjusted to facilitate distinguishing nuclear signal. The mitral layer was demarcated, and then a blinded investigator scored each image by counting the total number of nuclei in the mitral layer and scoring them for the presence of c-Fos. Nuclei in the mitral layer were considered positive for c-Fos expression if the euchromatin of a nucleus was occluded by c-Fos expression on the same focal plane as the nucleus. The FIJI counting plugin was used for counting c-Fos expressing nuclei in the mitral layer of the accessory olfactory bulb. The percent of activated neurons is the fraction of mitral cell nuclei that were identified as positive for c-Fos. Mixed models were used to analyze c-Fos counts, with a fixed effect of the interaction of genotype and condition and a random effect of subject. Post-hoc analysis compared the estimated marginal means of each condition within each genotype.

### Analysis of Behavioral Data

#### Attacks

Mice were labeled “attackers” if they attacked the intruder mouse during any of the three trials. Fisher’s exact test was used to analyze the proportion of attackers in each genotype.

#### Anogenital sniffing

Mixed models were used to analyze the duration of anogenital sniffing, with fixed effects of genotype and trial and a random effect of subject.

## Author contributions

Conceptualization: KM (Lead), NBG (Equal)

Data curation: NBG (lead), KM (equal)

Formal analysis: NBG (lead), KM (supporting), PK (Supporting), JD (Supporting) ZZ (Supporting)

Funding acquisition: KM

Investigation: NBG (Lead), PK (supporting), RE (supporting), HP (supporting), NY (supporting), ZZ (supporting)

Methodology: NBG (Lead), KM (Equal)

Project administration: KM (lead), NBG (supporting)

Resources: NBG (Lead), PK (Equal), KM (Supporting)

Supervision: KM (Lead), NBG (Supporting)

Validation: NBG (Lead), KM (Supporting)

Visualization: NBG (lead), KM (supporting)

Writing – original draft: NBG (lead), KM (Equal)

Writing – review & editing: NBG (lead), KM (Equal), JD (supporting)

## Acknowledgements

The authors thank Irina Ruzina, Mackenzie Leung, Melina Updale, and Sanjana Jethani for their help with the resident intruder experiments. We would also like to thank Madison Freestone for helping refine and edit the manuscript.

The Tex15 monoclonal antibody was produced by Susan Brenner-Morton, Director of the Antibody Platform within the Scientific Platforms at the Zuckerman Institute, Columbia University. We gratefully acknowledge her expertise in the development of this antibody. We are grateful for the support of the Antibody Platform and the Zuckerman Institute Scientific Platforms.

We acknowledge the School of Arts and Science-Human Genetics Institute of NJ (SAS-HGINJ) Imaging Core Facility for use of their confocal and widefield microscope.

## Data availability

Raw and processed RNA-seq data reported in this paper are available from GEO under accession number GSE326692. VNO scRNA-seq data were previously described (Hills et al., 2024) and are available from GEO under accession number GSE252365.

## Code availability

All code required to reproduce the figures in this manuscript is available at https://github.com/MonahanLab/Tex15_in_VNO/tree/main.

---

**Figure S1.**
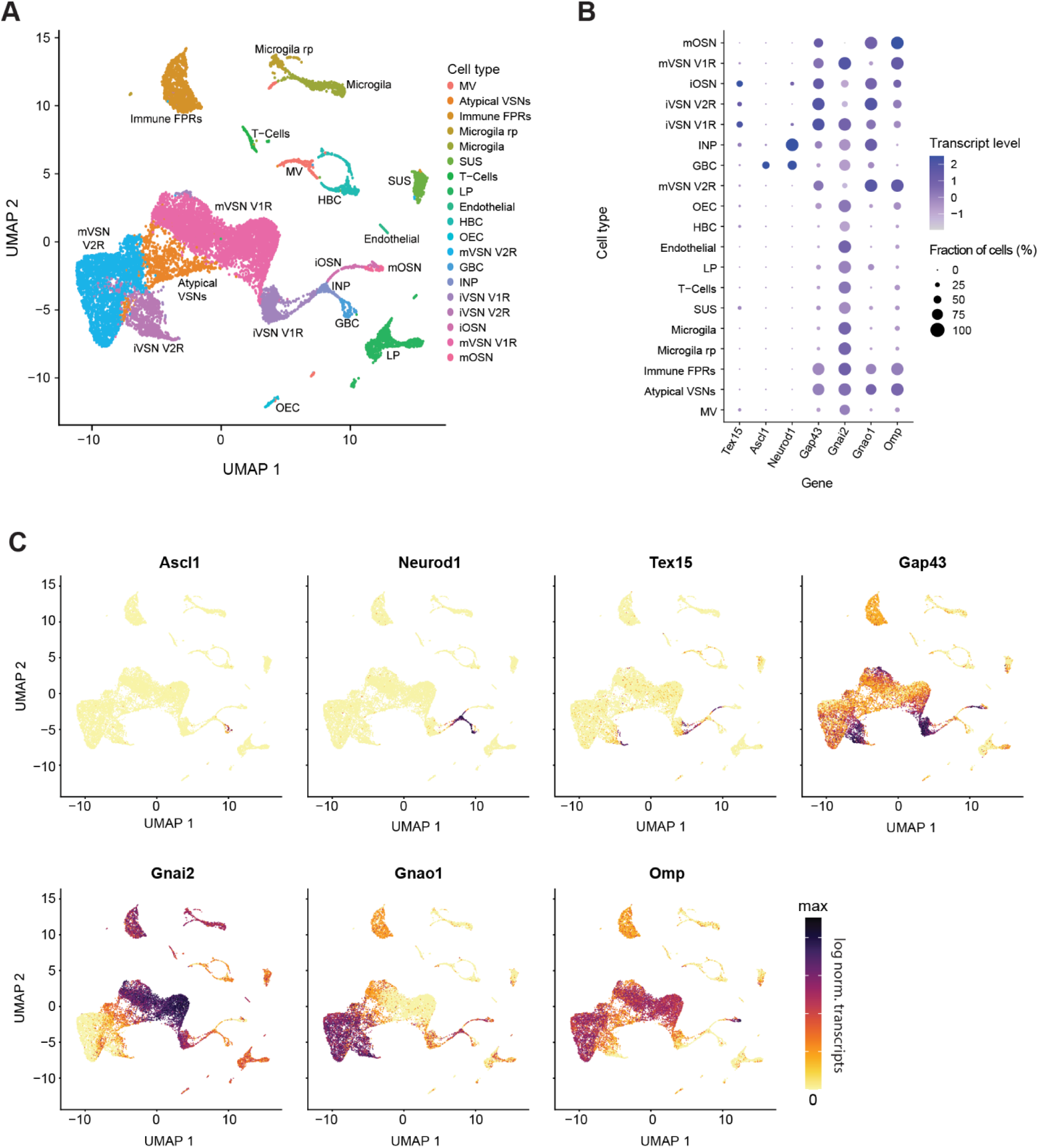
Identifying Tex15 expressing cells within the VNO. (A) UMAP projection of all cells present in VNO scRNA-seq from 14-day old mice. Abundant cell clusters are labeled based upon marker gene expression. (B) Dot plot showing transcript levels and frequency of expression for *Tex15* and VSN developmental markers across each identified cell cluster. (C) Transcript levels for developmental marker genes and Tex15 in all VNO cells. All gene expression UMAP and dot plots show transcript levels expressed as log-normalized counts. Atypical cells expressing both V1 and V2 VSN markers are called “Atypical VSNs”.

**Figure S2.**
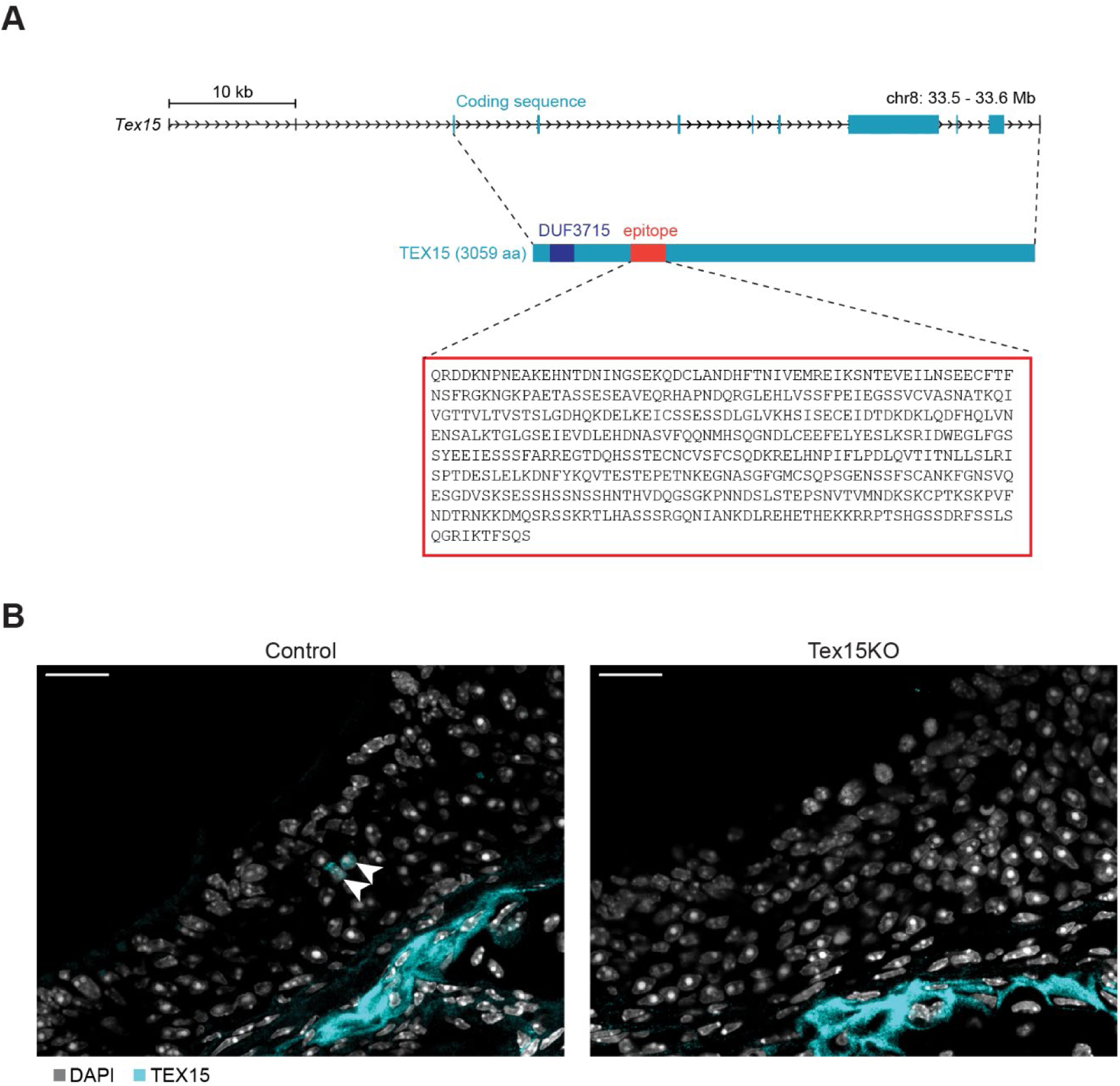
Detection of TEX15 expression. (A) Schematic showing the location and sequence used for generating a mouse monoclonal antibody against TEX15. (B) The Tex15 specific immunoreactivity (arrowheads) observed in VSN nuclei in VNO tissue sections from control mice is lost in Tex15KO VNO sections, while non-specific immunoreactivity typically observed with mouse monoclonal antibodies remains. TEX15 channel brightness was adjusted identically in both images. Scalebar: 20μm

**Figure S3.**
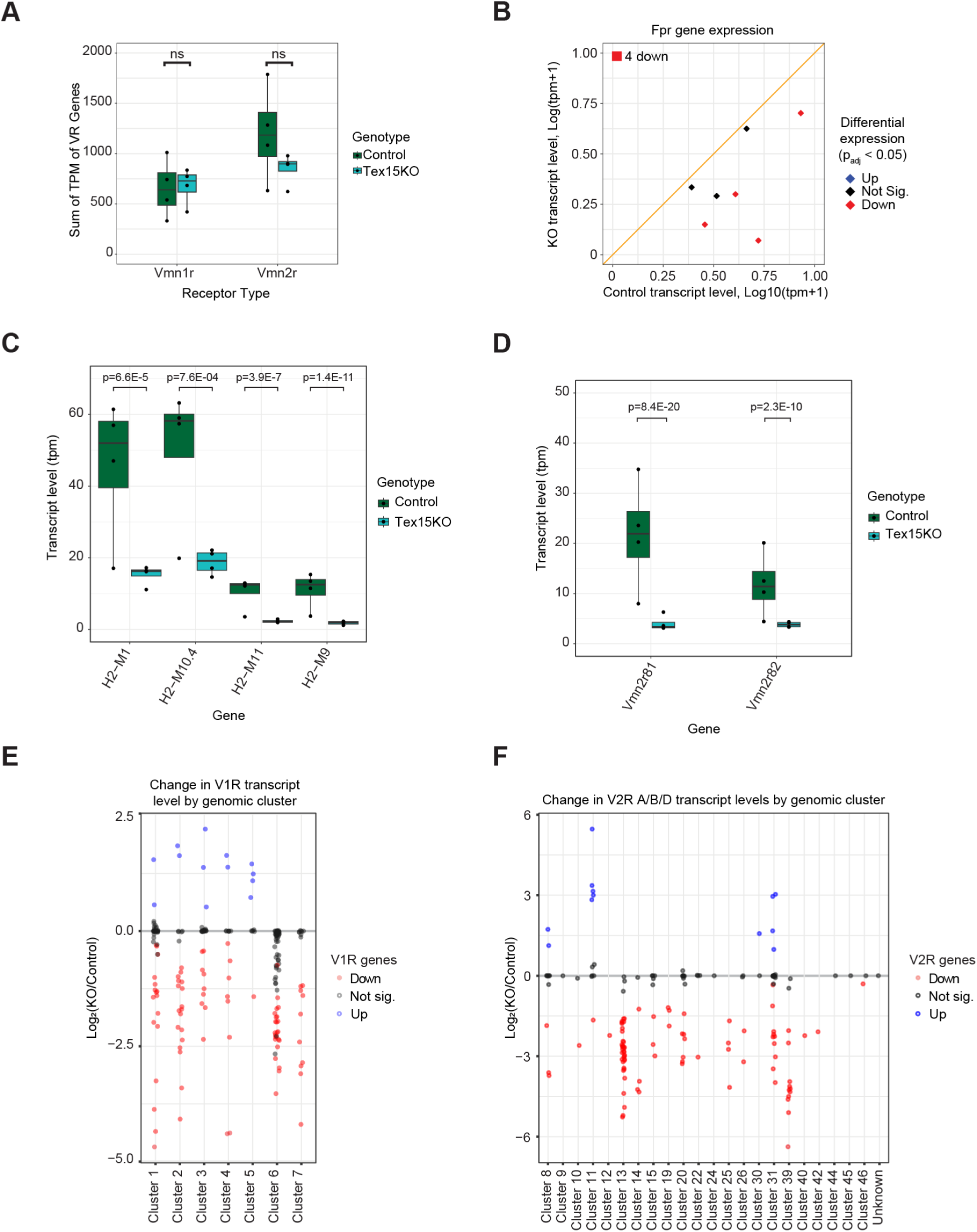
Transcriptional changes in the *Tex15* knockout. (A) Comparison of the total V1R and V2R transcript levels (sum of tpm) between the control and experimental conditions. T-test was conducted to determine significance. (B) Scatter plot of FPR transcript levels in Tex15KO versus control VNOs. Differentially expressed genes are in color. (C) Transcript levels of differentially expressed H2-MV genes. Significance was determined by a Wald Test on normalized counts. (D) Transcript levels of VR genes *Vmn2r81* and *Vmn2r82*, which are associated with the downregulated H2-MV genes. Significance was determined by a Wald Test on normalized counts. (E) Fold change of V1R transcript levels, with genes grouped by cluster. Differentially expressed genes are in color. (F) Fold change of A, B, and D V2Rs transcript levels with genes grouped by cluster. For bulk RNA-seq plots, differentially expressed genes were determined based upon padj < 0.05 for a >50% change in expression (Wald test). Genes that are significantly upregulated in the KO are blue, genes that are downregulated are red.

**Figure S4.**
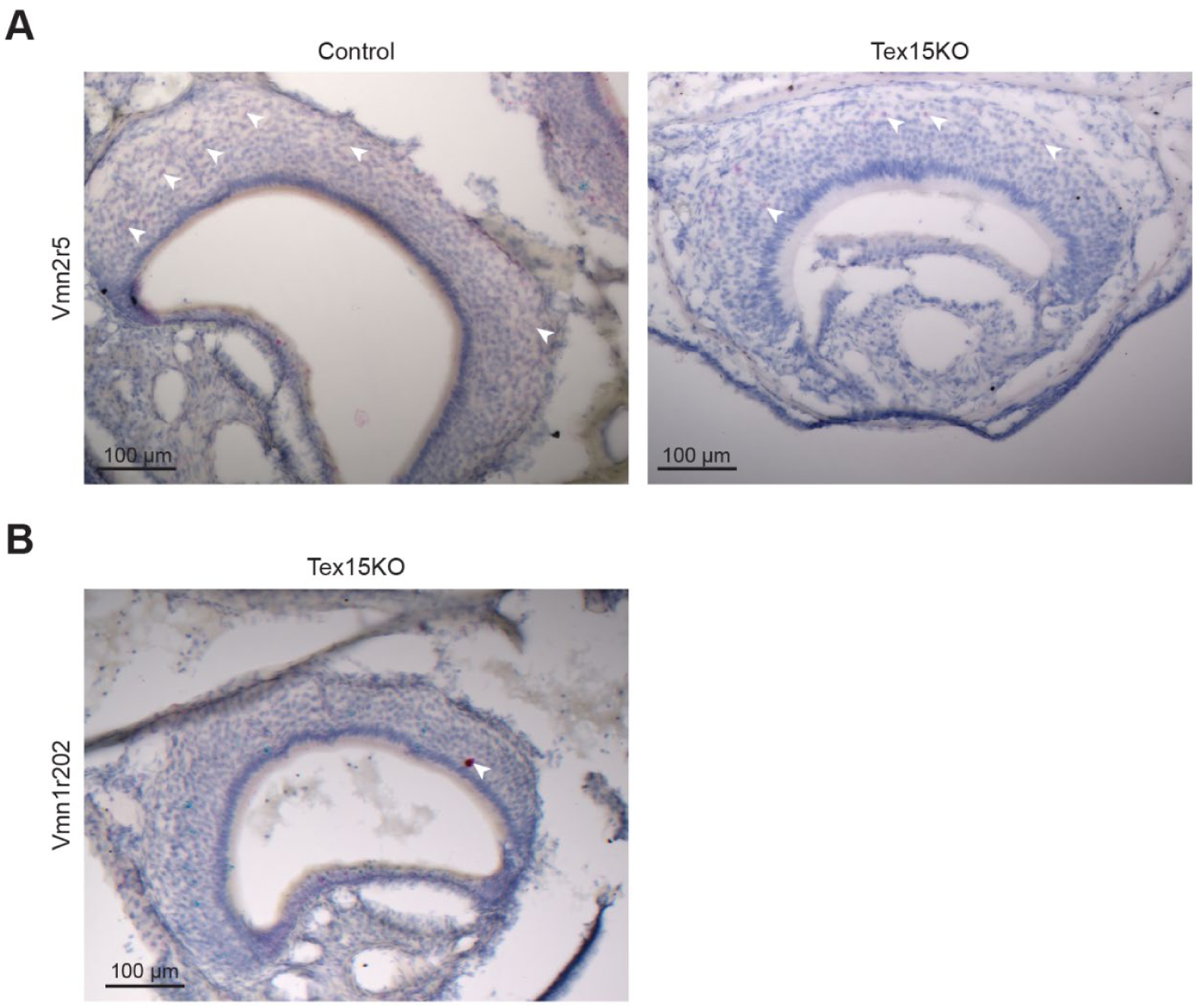
Tex15KO mice still express the downregulated VRs. (A) Representative images of *Vmn2r5* BaseScope assay on tissue sections from control and Tex15KO mice. Scalebar: 100μm (B) Representative image showing a Tex15KO VNO with a cell strongly expressing *Vmn1r202*. Scalebar: 100 μm.

